# Protein phosphatase 1 regulatory inhibitor subunit 14C promotes triple-negative breast cancer progression via sustaining inactive glycogen synthase kinase 3 beta

**DOI:** 10.1101/2021.05.09.443305

**Authors:** Yunting Jian, Lingzhi Kong, Hongyi Xu, Xinjian Huang, Yue Li, Dongni Shi, Yunyun Xiao, Muwen Yang, Siqi Li, Xiangfu Chen, Ying Ouyang, Yameng Hu, Xin Chen, Pian Liu, Weidong Wei

**Affiliations:** Department of Experimental Research, Sun Yat-sen University Cancer Center, State Key Laboratory of Oncology in South China, Collaborative Innovation Center for Cancer Medicine, Guangzhou, 510060, China.; Department of Breast Surgery, Sun Yat-Sen University Cancer Center, State Key Laboratory of Oncology in South China, Collaborative Innovation Center for Cancer Medicine, Guangzhou, 510060, China.; Department of Biochemistry, Zhongshan School of Medicine, Sun Yat-sen University, Guangzhou, 510080, China.; Departments of Pathophysiology, School of Basic Medical Sciences, Guangzhou Medical University, Guangzhou, 511436, China.; Cancer Center, Union Hospital, Tongji Medical College, Huazhong University of Science and Technology, Wuhan, 430022, China.

**Keywords:** TNBC, PPP1R14C, GSK3β, phosphorylation, degradation

## Abstract

The majority of clinical deaths among patients with triple-negative breast cancer (TNBC) are caused by uncontrolled cell proliferation and aggressive metastases, which is regulated by hyperactive glycogen synthase kinase 3 beta (GSK3β); however, the underlying mechanisms remain largely unknown. In the present study, we found that protein phosphatase 1 regulatory inhibitor subunit 14C (PPP1R14C) was specifically upregulated in TNBC, compared with that in normal tissues and non-TNBC. High PPP1R14C expression correlated significantly with shorter 5-year relapse-free survival and overall survival in patients with TNBC. Overexpressing PPP1R14C promoted, while suppression of PPP1R14C decreased the cell proliferation and the aggressive phenotype of TNBC cells, *in vitro* and *in vivo*. Importantly, we revealed that PPP1R14C interacted with and inactivated type 1 Ser/Thr protein phosphatase (PP1) to sustain GSK3β phosphorylation at S9, and induced the ubiquitylation and degradation of non-phosphorylated GSK3β (S9A) via recruiting E3 ligase, TRIM25. Furthermore, treating with C2 ceramide (C2), which recovered kinase activity of GSK3β, resulted in tumor growth inhibition in PPP1R14C-overexpressing TNBC cells. Our findings reveal a novel mechanism whereby PPP1R14C sustains inactive GSK3β, which might lead to targeted therapy for TNBC.

## Introduction

Triple-negative breast cancer (TNBC) is an aggressive breast cancer subtype characterized by the absence of the estrogen receptor (ER), progesterone receptor (PR) and human epidermal growth factor-2 (HER-2) amplification [1]. Lack of the receptors listed above means that TNBC does not respond to endocrine therapy or anti-HER-2 therapies; therefore, treatment is usually based on traditional chemotherapy [2, 3]. Uncontrolled proliferation and rapid recurrence are the main characteristics and challenges in treating TNBC, which might reduce the efficacy of chemotherapy [4]. Hence, there is an urgent need to develop new and effective treatments for TNBC, especially those involving actionable molecular targets.

Glycogen synthase kinase 3 beta (GSK3β), a highly evolutionarily conserved Ser/Thr kinase, is involved in diverse physiological and pathological processes via its selective phosphorylation of many substrates [5–7]. GSK3β is constitutively active in basal-state cells and acts as a tumor suppressor by phosphorylating oncogenic transcription factors (TFs), leading to their degradation [8]. When activated by oncogenic pathways, GSK3β is phosphorylated into an inactive state at Ser9, which leads to the accumulation and activation of downstream oncogenic signaling [9–13]. Elevated phosphorylated p-GSK3β (Ser9, inactive form) promoted epithelial-mesenchymal transition and metastasis in non-small cell lung carcinoma cells by accumulating Slug and Snail proteins [14]. Furthermore, decreased p-GSK3β (Ser9) levels inhibited cell proliferation and 7,12-dimethylbenz[a]anthracene (DMBA)-induced tumor growth in TNBC cell lines [15]. Therefore, p-GSK3β (Ser9) might play an important role in the progress of neoplastic diseases. The main phosphatase that reverses the phosphorylation state of GSK3β is protein phosphatase like type 1 Ser/Thr protein phosphatase (PP1), which is the key regulator in keeping the dynamically balance [16]. However, studies observed that GSK3β is an inactive form in human cancers, including TNBC, indicating widespread compromised phosphatase activity in tumors [17–19]. However, the molecular mechanisms in regulating GSK3β in TNBC remain unknown.

*PPP1R14C* has been discovered among morphine-regulated brain mRNAs, and encodes protein phosphatase 1 regulatory inhibitor subunit 14C, which has an inhibitory effect on PP1 [20, 21]. Daniel Horvath et al have revealed that PPP1R14C decreases the release of neurotransmitters by sustaining synaptosome associated protein 25 (SNAP25) in the phosphorylated state to regulate neuronal exocytosis [22]. Moreover, overexpression of PPP1R14C increased the phosphorylated RB transcriptional corepressor 1 (RB1), promoting leukemic cell survival under chemotherapy treatment [23]. Thus, PPP1R14C might be involved in human cancer progression by regulating the protein phosphorylation state. However, the functions and mechanism of PPP1R14C in TNBC remains to be explored.

In the present study, we found that PPP1R14C was specifically overexpressed in TNBC compared with that in normal tissues and non-TNBC and demonstrated that PPP1R14C played a vital role in promoting tumorigenesis of TNBC by sustaining a steady-state of p-GSK3β (S9) via compromising the ability of PP1 to dephosphorylate GSK3β, and inducing the ubiquitylation and degradation of non-phosphorylated GSK3β (S9A), which specifically recruited E3 ligase, TRIM25. Importantly, C2 ceramide (C2) treatment reversed the malignant phenotype induced by PPP1R14C, indicating a potential novel strategy to treat TNBC.

## Results

### PPP1R14C is specifically upregulated in TNBC

The phosphorylation state of GSK3β is balanced by kinases and phosphatases; however, the underlying mechanism for the persistent inhibition state of GSK3β in TNBC is unknown. To explore the mechanism of GSK3β regulation in TNBC, we first analyzed public cancer datasets from The Cancer Genome Atlas (TCGA) and identified 48 genes that were specifically upregulated in TNBC, compared with normal tissues and non-TNBC cancer tissues (Figure 1A and 1B). Among them, we focused on PPP1R14C because its inhibitory effect on PP1, the predominant phosphatase regulating GSK3β [20]. PPP1R14C mRNA levels were robustly upregulated in TNBC samples in the TCGA and Gene expression Omnibus (GEO) dataset (Figure 1C). Significantly, PPP1R14C was markedly upregulated in basal-like breast cancer (BLBC) tissues, followed by normal tissues and other subtypes (Luminal A, Luminal B and Her-2) (Figure 1D). Next, we detected PPP1R14C expression in breast cancer tissues and cell lines. Both the mRNA and protein levels of PPP1R14C were upregulated significantly in TNBC samples and cell lines compared with those in normal tissues and non-TNBCs (Figure 1E and 1F). Thus, these results suggested a substantial increase of PPP1R14C in TNBC.

**Figure 1.**
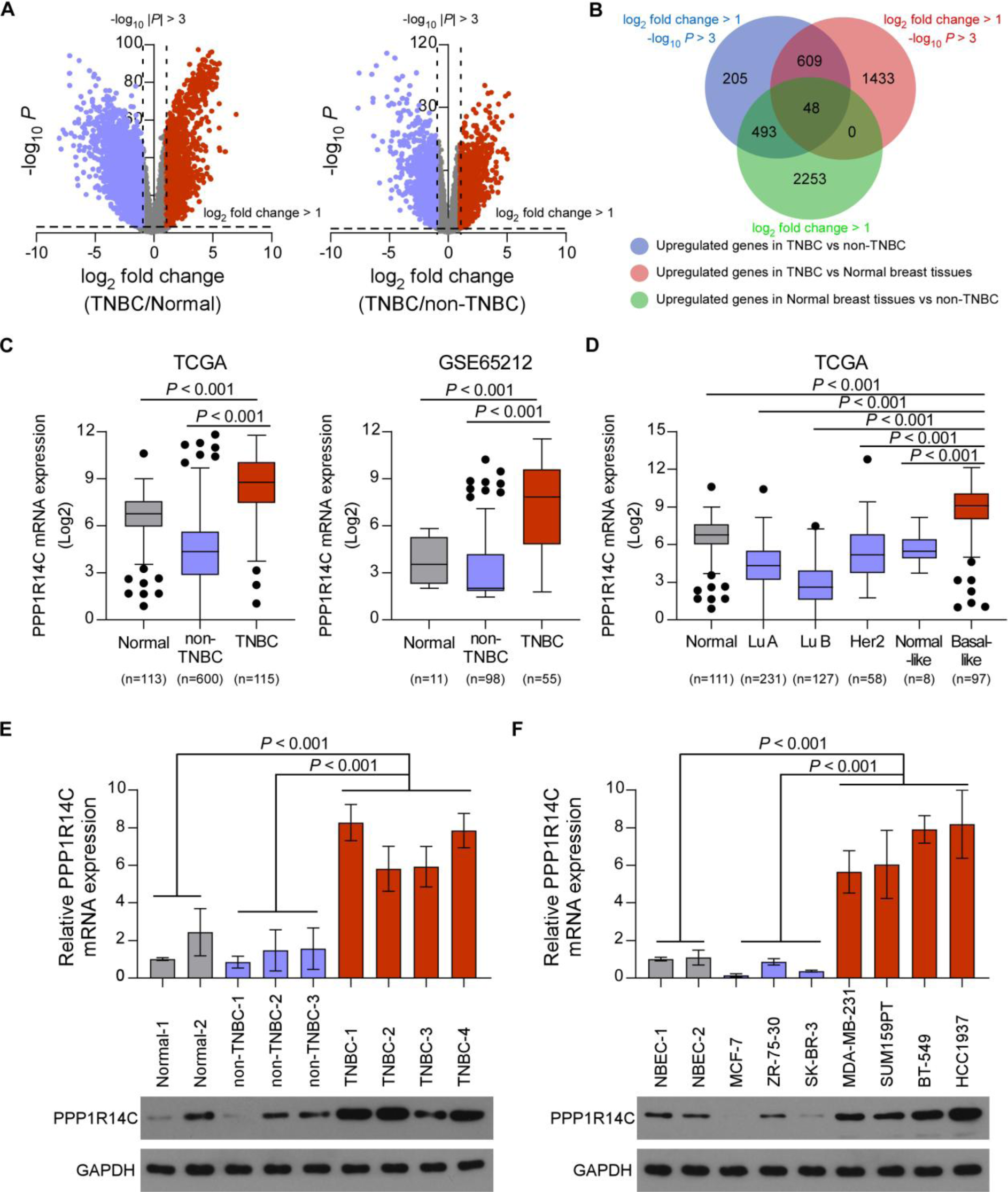
PPP1R14C is specifically upregulated in TNBC. **(A)** Volcano plot for gene expression in The Cancer Genome Atlas (TCGA) breast cancer dataset by comparing TNBC and normal tissues (left panel), and comparing TNBC and non-TNBC (right panel). The blue dots represent the downregulated genes and red dots represent the upregulated genes dysregulated in TNBC compared to normal samples/ non-TNBC. **(B)** A Venn diagram among upregulated genes in TNBC compared to non-TNBC, TNBC compared to normal samples, and normal tissues compared to non-TNBC. **(C)** PPP1R14C mRNA expression levels in The Cancer Genome Atlas (TCGA) breast cancer dataset (including 113 normal, 115 TNBC, and 600 non-TNBC samples) and Gene expression Omnibus (GEO) dataset (GSE65212, including 11 normal, 55 TNBC, and 98 non-TNBC samples). **(D)** PPP1R14C mRNA expression levels in tumor samples with informed molecular subtypes from TCGA breast cancer dataset. Statistic analysis was normalized to the expression levels in basal-like subgroup. Lu A, luminal A; Lu B, luminal B. **(E and F)** Real-time PCR (up) and western blot (down) analysis of PPP1R14C expression in human breast cancer tissues (E) and cell lines (F). GAPDH was used as a loading control. Data represent the means ± S.D. of three independent experiments.

### High expression of PPP1R14C is associated with poor prognosis in patients with TNBC

The clinical significance of high PPP1R14C expression was further assessed by immunohistochemistry (IHC) staining in 10 adjacent normal tissues and 150 archived breast cancer tissue samples, including 50 non-TNBC cases and 100 TNBC cases. IHC analysis revealed that PPP1R14C was upregulated markedly in TNBC, but was only detectable at low levels in normal breast tissues and non-TNBC tissues. The specimens with a staining index (SI) ≥ 6 were defined as PPP1R14C-high, while the others were PPP1R14C-low (Figure 2A). Correlation analysis showed that high PPP1R14C expression was associated significantly with advanced T classification (*P* = 0.013), and relapse status within 5 years (*P* = 0.002) in patients with TNBC (Figure 2B, Supplementary Table 2). Importantly, Kaplan-Meier survival curves and log-rank tests revealed that patients with high PPP1R14C expression suffered significantly poorer 5-year overall survival (OS) and relapse-free survival (RFS) in the TNBC subgroup (*P* = 0.004, hazard ratio (HR) = 5.046, 95% confidence interval (CI) = 2.097-12.14; *P* < 0.001, hazard ratio (HR) = 6.164, 95% confidence interval (CI) = 2.72-13.97, respectively); Figure 2C, Supplementary Table S3). In addition, the available online database Kaplan-Meier Plotter (http://kmplot.com/analysis) showed that breast cancer patients with high PPP1R14C expression had significantly shorter RFS in all breast cancer patients and basal-like subgroup (Figure 2D). Moreover, multivariate Cox regression analysis indicated that high PPP1R14C expression and T classification as independent prognostic factors for 5-year OS and 5-year RFS in TNBC (Figure 2E and Supplementary Table S3). Thus, upregulation of PPP1R14C might contribute to TNBC progression, leading to a poor clinical outcome.

**Figure 2.**
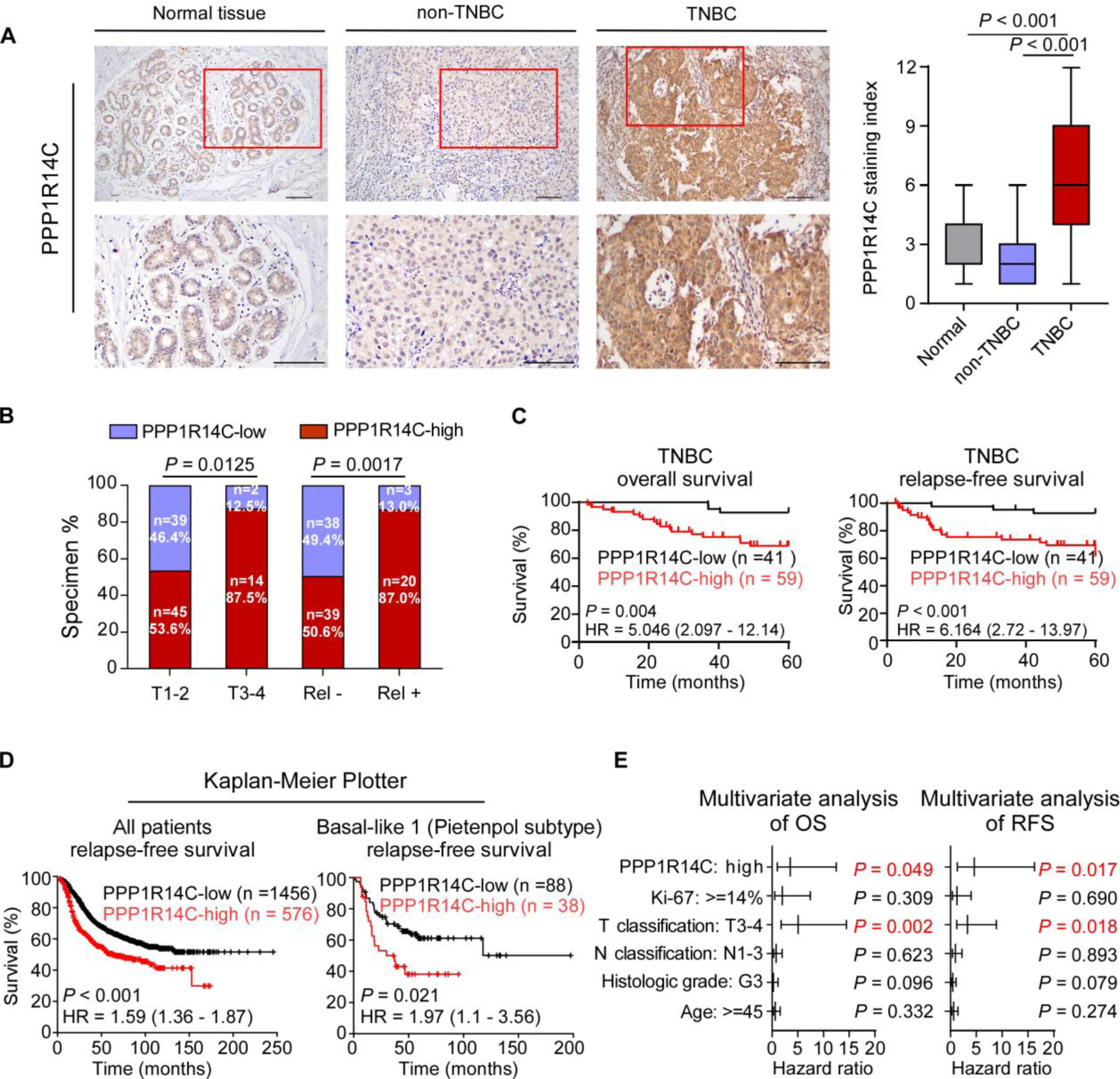
Upregulation of PPP1R14C is associated with poor prognosis in TNBC. **(A)** Representative images of PPP1R14C staining in normal breast tissues, non-TNBC and TNBC tissues. The SI distribution of each group was indicated (right panel) and Mann-Whitney U test was used for analysis. Scale bars represent 100 μm. **(B)** The distribution and correlation between PPP1R14C staining and T classification and relapse status in patients with TNBC. The χ^2^ test was used for analysis. Rel -, no relapse; Rel +, with relapse. **(C)** Kaplan-Meier 5-year overall survival and relapse-free survival curve for patients with TNBC stratified by low (n = 41) and high PPP1R14C expression (n = 59, log-rank test). HR, hazard ratio. **(D)** The Kaplan-Meier Plotter (http://kmplot.com/analysis) program was used to analyze relapse-free survival of all breast cancer patients and basal-like subgroup (Pietenpol subtype). All settings were left at default values except for: gene symbol (PPP1R14C), survival (RFS), and auto select best cutoff (on). **(E)** Multivariate Cox regression analysis to evaluate the significance of the association between PPP1R14C signature, and OS and RFS in the presence of other clinical variables.

### PPP1R14C promotes tumor progression in TNBC cells *in vitro*

To further investigate the biological role of PPP1R14C in TNBC progression, we first stably overexpressed PPP1R14C or silenced endogenous PPP1R14C expression in MDA-MB-231 and SUM159PT breast cancer cell lines (Figure 3A). Strikingly, PPP1R14C-overexpressing TNBC cells displayed higher growth rates and increased anchorage-independent growth ability compared with vector-control cells, and had increased proportions of cells in the S phase and reduced proportions of cells in the G1 phase (Figures 3B-3E, Supplementary Figure 1A and 1B). In addition, upregulation of PPP1R14C strongly increased the mobility and invasion of TNBC cells (Figures 3F and 3G, Supplementary Figure 1C and 1D). Correspondingly, silencing PPP1R14C reduced the growth rates and anchorage-independent growth ability significantly, decreased the proportions of cells in the S phase and increased proportions of cells in the G1 phase, and decreased the TNBC cells invasion (Figure 3B-3G and Supplementary Figure 1A-1D). Collectively, these results suggested that PPP1R14C plays an important role of TNBC aggressiveness.

**Figure 3.**
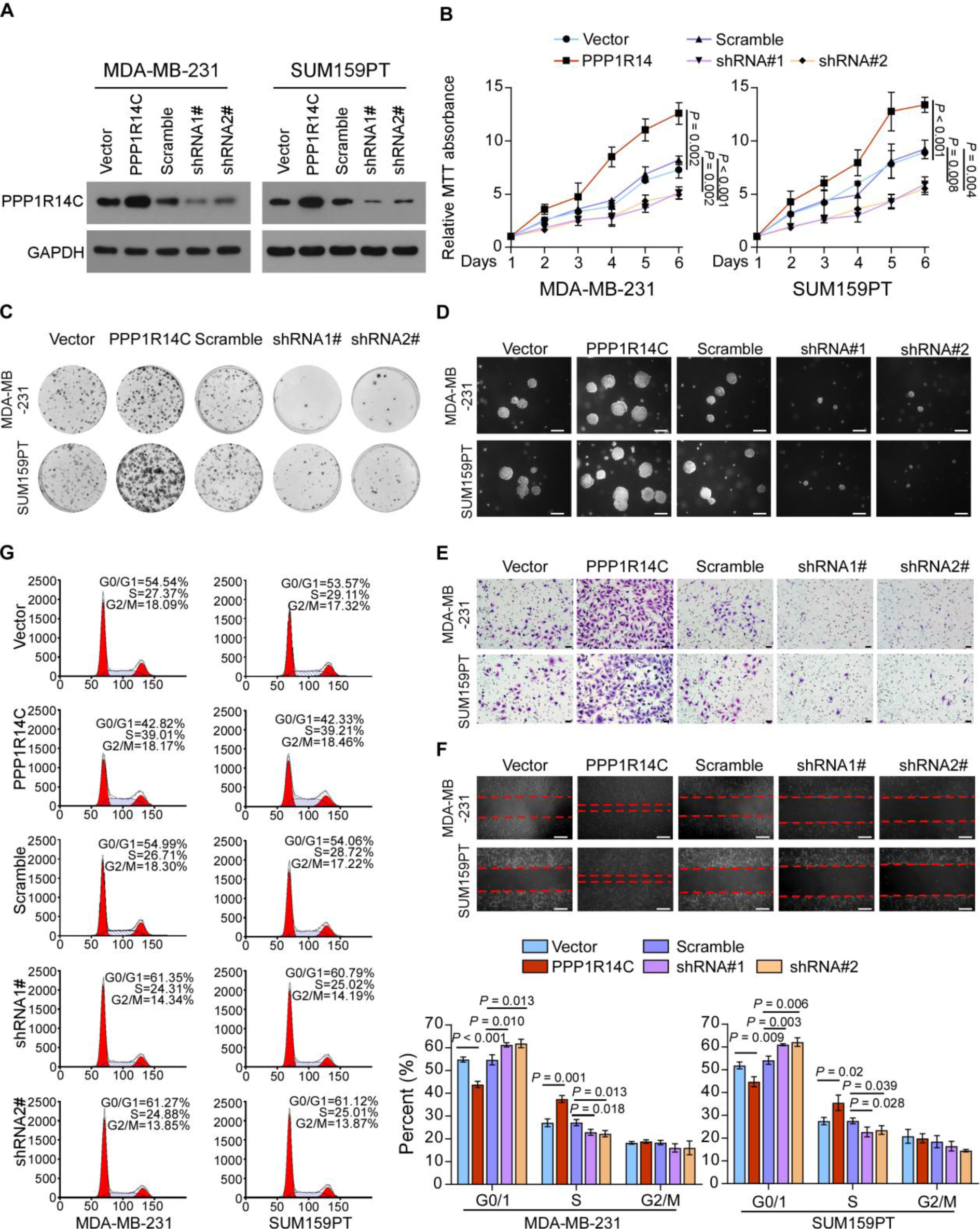
PPP1R14C promotes tumor progression in TNBC cells *in vitro*. **(A)** Western blotting analysis of PPP1R14C in MDA-MB-231 and SUM159PT cells stably transduced with PPP1R14C-overexpressing and PPP1R14C-silencing plasmids. GAPDH was used as a loading control. **(B-G)** MTT (B), colony formation (C), soft agar (D), transwell (E), wound healing (F) assays and flow cytometric analysis (G) were performed in the indicated cells. Scale bars represent 50 μm. Two-tailed Student’s t test was used. Data represent the means ± S.D. of three independent experiments.

### PPP1R14C contributes to TNBC tumorigenesis and metastasis

Next, we assessed the effect of PPP1R14C on tumorigenesis *in vivo*. We generated xenografts through subcutaneous injection of stable SUM159PT cell lines and measured tumor growth. Compared with the control groups, tumor growth was markedly enhanced in the PPP1R14C-overexpressing group and suppressed in PPP1R14C silenced group (Figure 4A and Supplementary Figure 2A). Tumors overexpressing PPP1R14C had high expression of the proliferation marker Ki-67 compared with that in the vector group, whereas the PPP1R14C-silenced tumors showed decreased Ki-67 expression (Figure 4B and Supplementary Figure 2B). Next, the effects of PPP1R14C on breast cancer metastasis were examined in lung colonization models. Stable SUM159PT cell lines were injected into Balb/c nude mice via the lateral tail veins. Metastatic activity was evaluated using bioluminescence imaging (BLI) of luciferase-transduced cells and by hematoxylin and eosin (H&E) staining of lung metastases sections. Upregulation of PPP1R14C increased the lung metastasis burden (Figure 4C). Importantly, overexpressing PPP1R14C decreased the survival time and proportion of mice strikingly (Figure 4D). In contrast, downregulation of PPP1R14C significantly reduced lung tumor outgrowth and mouse deaths (Figure 4C and 4D).

**Figure 4.**
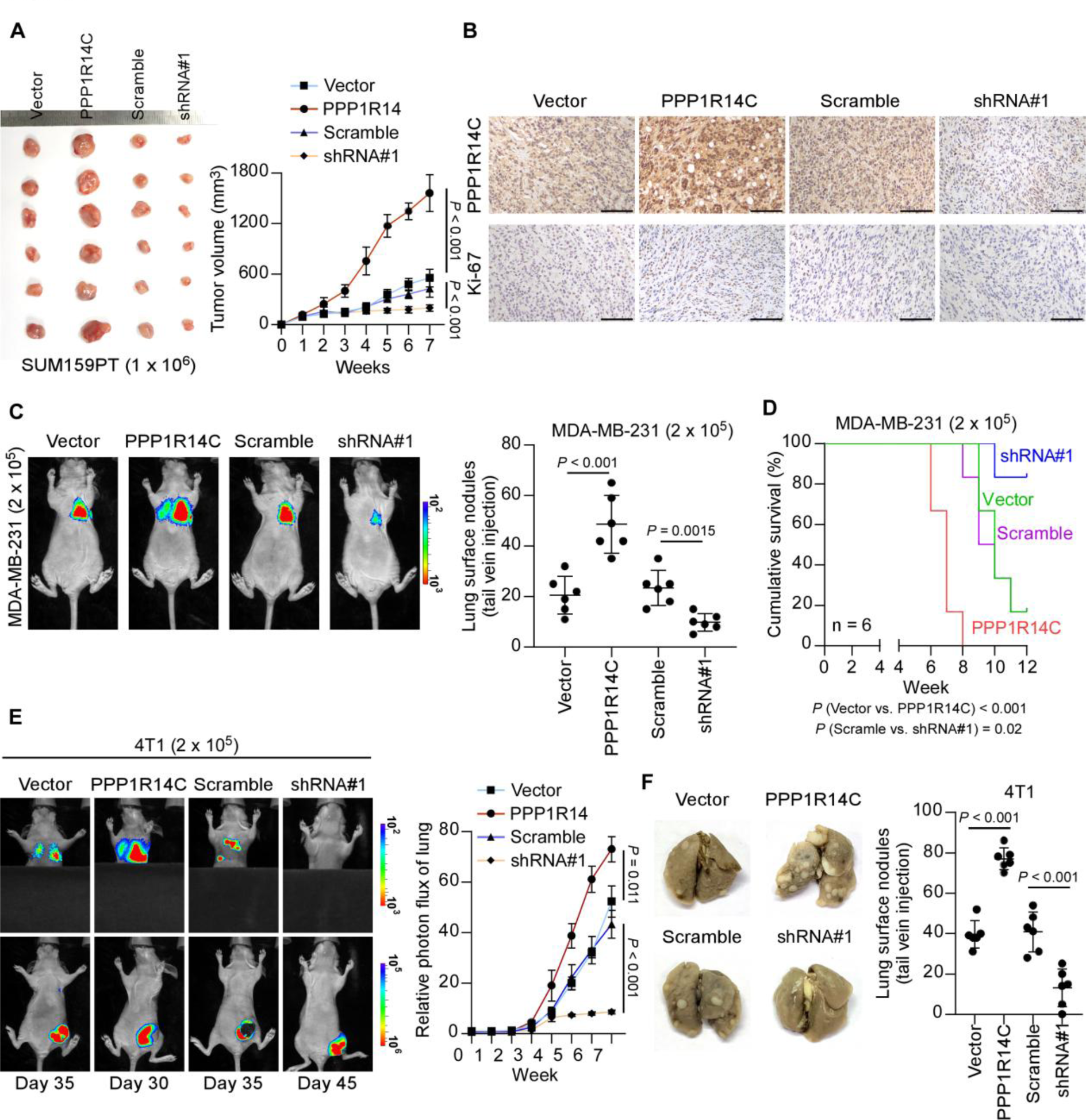
PPP1R14C contributes to TNBC tumorigenesis and metastasis. **(A)** PPP1R14C-overexpressing, PPP1R14C-silenced and vector/scramble SUM159PT cell lines were subcutaneously injected into mice (1 × 10^6^/injection, n = 6/group). The tumor volumes in each group are shown. Data represent the means ± S.D. of three independent experiments. **(B)** IHC of Ki-67 staining showed in the indicated xenografts. Scale bars represent 100 μm. **(C)** *In vivo* metastasis assays of PPP1R14C-overexpressing, PPP1R14C-silenced and vector/scramble MDA-MB-231 cells. Lung metastasis burden of xenografted animals was monitored weekly using bioluminescent imaging (BLI). Representative BLI images of the lungs were shown. The visible surface metastatic lesions were counted. **(D)** Kaplan–Meier survival curves of mice. **(E)** Lung colonization and spontaneous metastasis models of PPP1R14C-overexpressing, PPP1R14C-silenced and vector/scramble 4T1 cells. BLI quantification of lung metastasis of the indicated cells. **(F)** The visible surface metastatic lesions of lungs were counted. Two-tailed Student’s t test and log-rank test were used.

We further examined the role of PPP1R14C in a spontaneous metastatic model. Mice were injected subcutaneously with the luciferase-expressing 4T1 cells (1 × 10^5^), which are highly metastatic. Strikingly, BLI revealed that the metastasis of 4T1 cells was promoted strongly in the PPP1R14C overexpression group, but suppressed in the PPP1R14C silenced group, which was confirmed by counting the visible metastatic lesions (Figure 4E and 4F). Moreover, upregulating PPP1R14C shortened the survival time of mice strikingly (Supplementary Figure 2C). These findings indicated that PPP1R14C enhances tumorigenesis and metastasis of TNBC.

### PPP1R14C interacts with PP1 to constantly inhibit GSK3β activity

PPP1R14C is a powerful inhibitor for PP1, a major phosphatase responsible for modulating the phospho- and dephospho-balance of numerous substrate proteins[24]. To identify whether GSK3β is the exact substrate protein affected by the PPP1R14C and PP1 interaction in TNBC cells, we screened a collection of PP1 substrates [25–30]. The results indicated that overexpressing PPP1R14C significantly increased, while silencing PPP1R14C decreased the level of phosphorylation GSK3β, but did not influence the other substrate proteins (Figure 5A). In addition, PPP1R14C only affected the level of phosphorylation GSK3β at Ser9, but not Tyr216, the other inactive site of GSK3 β (Figure 5A). Importantly, PPP1R14C overexpression decreased, while knockdown of PPP1R14C increased the expression levels of known GSK3β downstream genes (Figure 5B), indicating that PPP1R14C could sustain phosphorylated GSK3β. Reciprocal immunoprecipitation (IP) assays using antibodies against PPP1R14C and PP1 demonstrated that PPP1R14C/PP1/p-GSK3β (Ser9) formed a complex in SUM159PT cells (Figure 5C). IP assays using HA-GSK3β transduced into Hela cells together with Flag-PPP1R14C and myc-PP1 were performed under treatment of SHIP2-IN-1, a GSK3β inhibitor that deactivates GSK3β via phosphorylating at Ser9 [31]. The results showed that PPP1R14C and PP1 specifically interacted with p-GSK3β (Ser9) (Figure 5D). Additionally, silencing PP1 did not abrogated the interaction of PPP1R14C with p-GSK3β (Ser9) in TNBC cell lines, indicating that PPP1R14C binds to p-GSK3β (Ser9) in a PP1-independent manner endogenously (Figure 5E). Furthermore, PPP1R14C overexpression decreased PP1 activity, and increased the p-GSK3β level, while silencing PPP1R14C showed reversed effects (Figure 5F-5I). These results suggested that PPP1R14C interacts with PP1 and inhibits PP1-induced p-GSK3β (Ser9) dephosphorylation.

**Figure 5.**
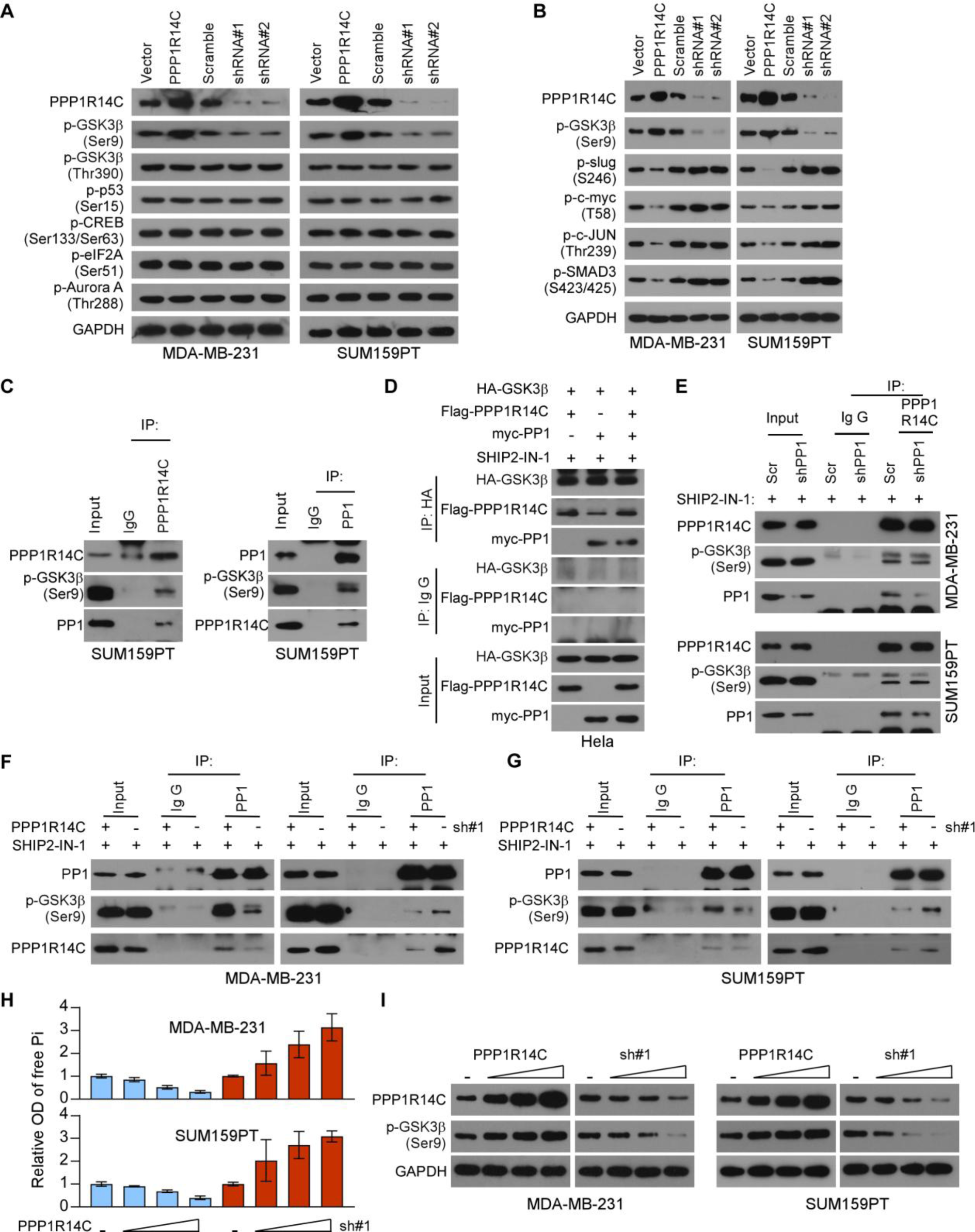
PPP1R14C interacts with PP1 to constantly inhibit GSK3β activity. **(A)** Western blot analysis of PPP1R14C, p-GSK3β (Ser9), p-p53 (Ser15), p-CREB (Ser133/Ser63), p-eIF2A (Ser51), and p-Aurora A (Thr288) in PPP1R14C-transduced cells and PPP1R14C-silenced cells compared with control cells. GAPDH was used as a loading control. **(B)** Western blot analysis of PPP1R14C, p-GSK3β (Ser9), p-slug (S246), p-c-myc (T58), p-c-Jun (Thr239), and p-SMAD3 (S423/425) in PPP1R14C-transduced cells and PPP1R14C-silenced cells compared with control cells. GAPDH was used as a loading control. **(C)** Reciprocal immunoprecipitation (IP) assay revealed the interaction of PPP1R14C, PP1 and p-GSK3β (Ser9) in SUM159PT cells. **(D)** Extraneous IP assay showed that the specifically interaction of PPP1R14C/PP1 with p-GSK3β (Ser9) under SHIP2-IN-1 treatment (10 µM, 24 hours). **(E)** Endogenous IP assay showed that PPP1R14C bound to p-GSK3β (Ser9) in a PP1-independent manner. **(F-G)** IP assays performed in TNBC cells treated under SHIP2-IN-1 with and without PPP1R14C-overexpressed or -silenced were subjected to immunoprecipitation with an anti-PP1 antibody followed by western blotting with the indicated antibodies. **(H)** The phosphatase activity of PP1 was measured by detecting its product, free Pi in the TNBC cells transfected with increased dose of PPP1R14C-expressing or PPP1R14C-shRNA plasmids. **(I)** The expression of p-GSK3β (Ser9) was detected by western blot assay in the TNBCs transfected with increased dose of PPP1R14C-expressing or PPP1R14C-shRNA plasmids. GAPDH was used as a loading control. The number of technical and biological replicates performed for the above assays depicted was three.

### PPP1R14C accelerates non-phosphorylated GSK3β (S9A) degradation

Interestingly, PPP1R14C overexpression increased p-GSK3β (Ser9) expression, but reduced the protein level of GSK3β, while did not alter its mRNA expression in TNBC cells (Figure 6A and Supplementary Figure 3A). It has reported that degradation of GSK3β is largely dependent on proteasome pathway in lung epithelial cells [32]. To determine whether PPP1R14C regulates the GSK3β stability, we detected its protein levels in the presence of cycloheximide (CHX) and MG132. Interestingly, the half-life of GSK3β was shortened remarkably under these treatments in PPP1R14C-overexpressing SUM159PT and MDA-MB-231 cells compared with that in the vector group, and the opposite effects were observed in the PPP1R14C-silenced cells (Figure 6B and Supplementary Figure 3B-3C).

**Figure 6.**
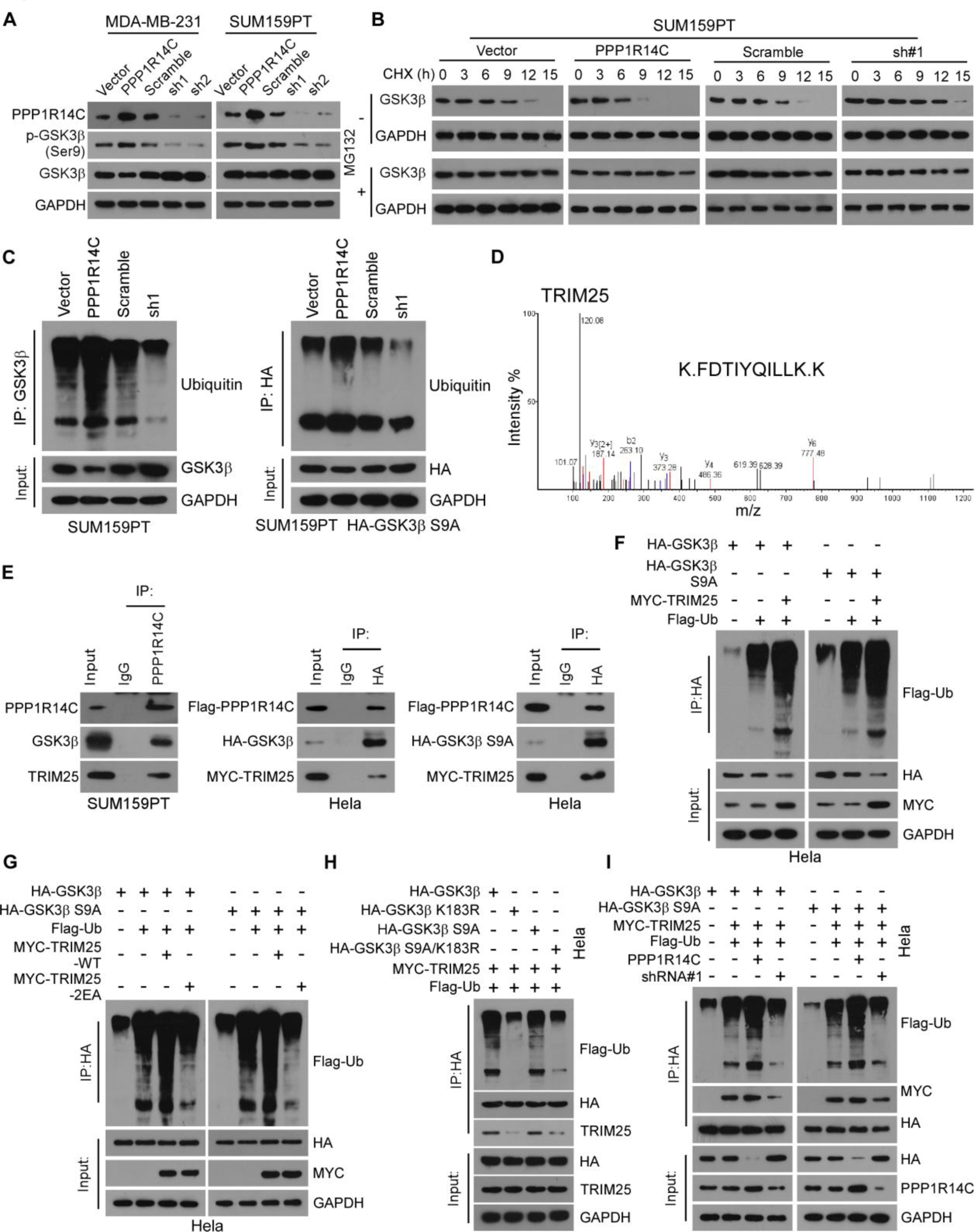
PPP1R14C accelerates non-phosphorylated GSK3β (S9A) degradation. **(A)** Western blot analysis of PPP1R14C, p-GSK3β (Ser9), and GSK3β in PPP1R14C-overexpressed cells and PPP1R14C-silenced cells compared with control cells. GAPDH was used as a loading control. **(B)** Western blot analysis of GSK3β protein in SUM159PT cells treated with CHX (50 µg/mL) for 0, 30, 60, or 120 min plus with or without MG132 (10 µM) treatment. GAPDH was used as loading control. **(C)** Effect of ubiquitination on GSK3β and non-phosphorylated GSK3β (S9A) by immunoprecipitation assay in SUM159PT or SUM159PT-HA-GSK3β (S9A) cells using anti-GSK3β or anti-HA antibody to pull down, respectively. **(D)** Representative MS plots and sequences of peptides from TRIM25. **(E)** Immunoprecipitation (IP) assay revealed the interaction between PPP1R14C, TRIM25 and GSK3β/ non-phosphorylated GSK3β (S9A) in SUM159PT and Hela cells. **(F)** Hela cells transfected with HA-GSK3β, non-phosphorylated HA-GSK3β (S9A) and Flag-Ub along with MYC-TRIM25 plasmids were subjected to immunoprecipitation with an anti-HA antibody followed by western blotting with an anti-Flag antibody. **(G)** Hela cells transfected with the plasmids as indicated were subjected to IP assay with an anti-HA antibody followed by western blotting with an anti-Flag antibody. **(H-I)** Hela cells transfected with the plasmids as indicated were subjected to IP assay with an anti-HA antibody followed by western blotting with the indicated antibodies. The number of technical and biological replicates performed for the above assays depicted was three.

Furthermore, altered PPP1R14C did not affect the half-life of p-GSK3β (Ser9) (Supplementary Figure 3D). In addition, we constructed TNBC cell lines stably expressing mutant HA-GSK3β (S9A), which fails to be phosphorylated. IP assays performed in SUM159PT, SUM159PT-HA-GSK3β (S9A) and Hela cell lines to detect the ubiquitination of different GSK3β forms. High levels of PPP1R14C promoted total and non-phosphorylated GSK3β (S9A) ubiquitination and low PPP1R14C levels inhibited this process, while p-GSK3β (Ser9) ubiquitination was not affected (Figure 6C and Supplementary Figure 4A-4B). The similar results were observed in MDA-MB-231 and MDA-MB-231-HA-GSK3β (S9A) cell lines (Supplementary Figure 4C).

To identify which E3 ubiquitin ligases is mediated by PPP1R14C to degrade non-phosphorylated GSK3β, mass spectrometry (MS) was used in SUM159PT. As shown in Figure 6D, tripartite motif containing 25 (TRIM25) was identified as a potent E3 ligase. Importantly, co-immunoprecipitation (Co-IP) using an anti-PPP1R14C antibody revealed that PPP1R14C could form complex with TRIM25 and GSK3β in SUM159PT cells. Furthermore, Co-IP was performed in Hela transfected with Flag-PPP1R14C, MYC-TRIM25, HA-GSK3β or HA-GSK3β (S9A) and confirmed that PPP1R14C mediated TRIM25-total/non-phosphorylated GSK3β interaction (Figure 6E). Next, we generated a series of truncated mutants of GSK3β and TRIM25 to dissect functional domains of the GSK3β-TRIM25 interaction. IP assays showed that only those GSK3β truncations with the N1 domain could bound to TRIM25, and N1 mutant with S9A was also found in TRIM25-MYC immunoprecipitate, suggesting that the interaction between GSK3β and TRIM25 required the N-terminal domains (Supplementary Figure 4D). For TRIM25, we constructed four TRIM25 truncates mutants, containing RING, B-box, coiled-coil (CC) and PRY/SPRY domain. And the immunoprecipitation assay showed that only TRIM25 truncated mutant containing B-box domain was necessary for its interaction with GSK3β (Supplementary Figure 4D). As a ubiquitin E3 ligase, TRIM25 probably binds to and mediates the ubiquitination of GSK3β, resulting in GSK3β degradation. To test this hypothesis, Flag-Ub, TRIM25, HA-GSK3β or HA-GSK3β (S9A) were expressed into Hela cells and co-immunoprecipitated with anti-Flag antibody. As shown in Figure 6F, TRIM25 promoted the total GSK3β and non-phosphorylated GSK3β-S9A ubiquitination. To elucidate the role of TRIM25 as an E3 ligase, we constructed a TRIM25 mutant with Glu9 and Glu10 changed into Ala (termed as TRIM25-2EA), which lost its ubiquitination activity [30, 33]. IP data showed that exogenous expression of the wild type (WT) of TRIM25, other than TRIM25-2EA significantly enhanced the total GSK3β and non-phosphorylated GSK3β-S9A ubiquitination (Figure 6G). In addition, K183 of GSK3β has been recognized as the ubiquitination site, and exogenous co-IP further suggested that the mutation of GSK3β K183R prompted the disassembly of ubiquitin and GSK3β and impeded the interplay of TRIM25 and GSK3β (Figure 6H). Furthermore, PPP1R14C overexpression substantially increased the interaction of total/non-phosphorylated GSK3β and TRIM25, while PPP1R14C silencing inhibited this interaction (Figure 6I). Therefore, these results showed that PPP1R14C enhanced the degradation of non-phosphorylated GSK3β by increasing TRIM25-dependent ubiquitination.

### Blockage of the PPP1R14C/PP1/p-GSK3β axis results in growth inhibition of TNBC cells with high-PPP1R14C levels *in vitro* and *in vivo*

To validate that the PPP1R14C/PP1/p-GSK3β axis is involved in TNBC tumor progression, we established PP1-silenced (using shPP1, a short hairpin RNA targeting PP1) TNBC cell lines or treated cells with C2, a PP1 activator to reverse the inhibitory effect of PPP1R14C[34]. As expected, shPP1 and C2 administration impaired PPP1R14C-induced cell growth, invasion, G1-S transformation, and anchorage-independent growth in TNBC cells, suggesting that PP1 is essential for PPP1R14C-mediated GSK3β inactivation (Figure 7A-7E). Next, we injected SUM159PT cells stably overexpressing PPP1R14C into mice subcutaneously and observed the tumor growth in each experimental group. Tumors formed by shPP1- or C2-treated cells were much smaller than those in the control group (Figure 7F and Supplementary Figure 5A). IHC of Ki-67 in tumor specimens showed that silencing PP1 or treating with C2 reduced the Ki-67 staining intensity markedly in PPP1R14C overexpressing tumors compared with that in the vehicle groups (Figure 7G and Supplementary Figure 5B). Furthermore, silencing PP1 or C2 treatment decreased the lung metastasis burden and mouse death in PPP1R14C-upregulated SUM159PT and 4T1 cells (Figure 7H-7J, Supplementary Figure 5C). In addition, we generated subcutaneous xenografts, a lung colonization model and a spontaneous metastatic model using TNBC vector cells. The effect of tumor growth and metastatic ability treating with silencing PP1 and C2 in vector cells comparable to that in PPP1R14C-overexpressing TNBC cells, only slightly repressed tumor growth and metastatic ability (Supplementary Figure 5D-5L). Collectively, these results suggested that blockage of the PPP1R14C/PP1/p-GSK3β axis might have anti-TNBC effects in the presence of a high-PPP1R14C level.

**Figure 7.**
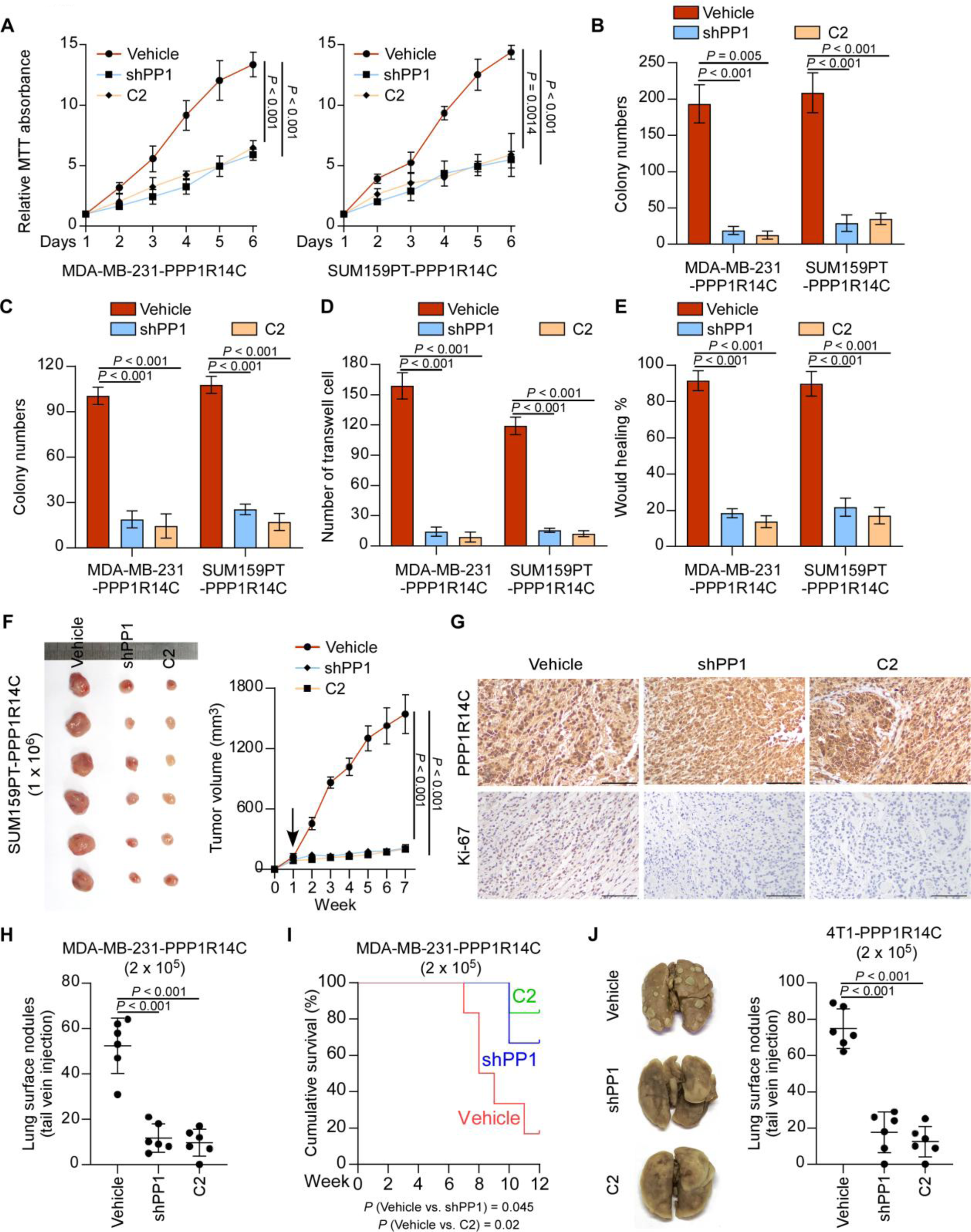
Blockage of the PPP1R14C/PP1/p-GSK3β axis results in growth inhibition of TNBC cells with high-PPP1R14C levels in vitro and in vivo. **(A-E)** MTT (A), colony formation (B), soft agar (C), transwell (D), and wound healing (E) assays were performed in vehicle, shPP1, or C2 (10 μM) in stably-overexpressed PPP1R14C MDA-MB-231 and SUM159PT cells. Data represent the means ± S.D. of three independent experiments. Two-tailed Student’s t test was used. **(F)** PPP1R14C-overexpressd cell lines SUM159PT with or without shPP1 were subcutaneously injected into mice (1 × 10^6^/injection, n = 6/group). One week after inoculation, control and shPP1 groups were treated with intraperitoneal 20%/ 80% DMSO/ saline, while treatment group received 100 mg/kg intraperitoneal C2 once per day. The tumor volumes in each group are shown. **(G)** IHC of PPP1R14C and Ki-67 staining showed in the indicated xenografts. Scale bars represent 100 μm. (H) *In vivo* metastasis assays of vehicle, shPP1 and C2 treatment groups. The visible surface metastatic lesions were counted. **(I)** Kaplan–Meier survival curves of mice. **(J)** The visible surface metastatic lesions of lungs were counted. Two-tailed Student’s t test and log-rank test were used.

### Clinical relevance of the PPP1R14C/PP1/p-GSK3β axis in TNBC

Finally, we assessed the PPP1R14C/PP1/p-GSK3β axis in clinical specimens. IHC staining for PP1 and p-GSK3β was performed in the same cohort of 100 TNBC patient specimens. PPP1R14C expression correlated strongly with PP1 and p-GSK3β (Ser9) levels, suggesting that the PPP1R14C/PP1/p-GSK3β axis was clinically relevant (Figure 8A and Supplementary Figure 6A).

**Figure 8.**
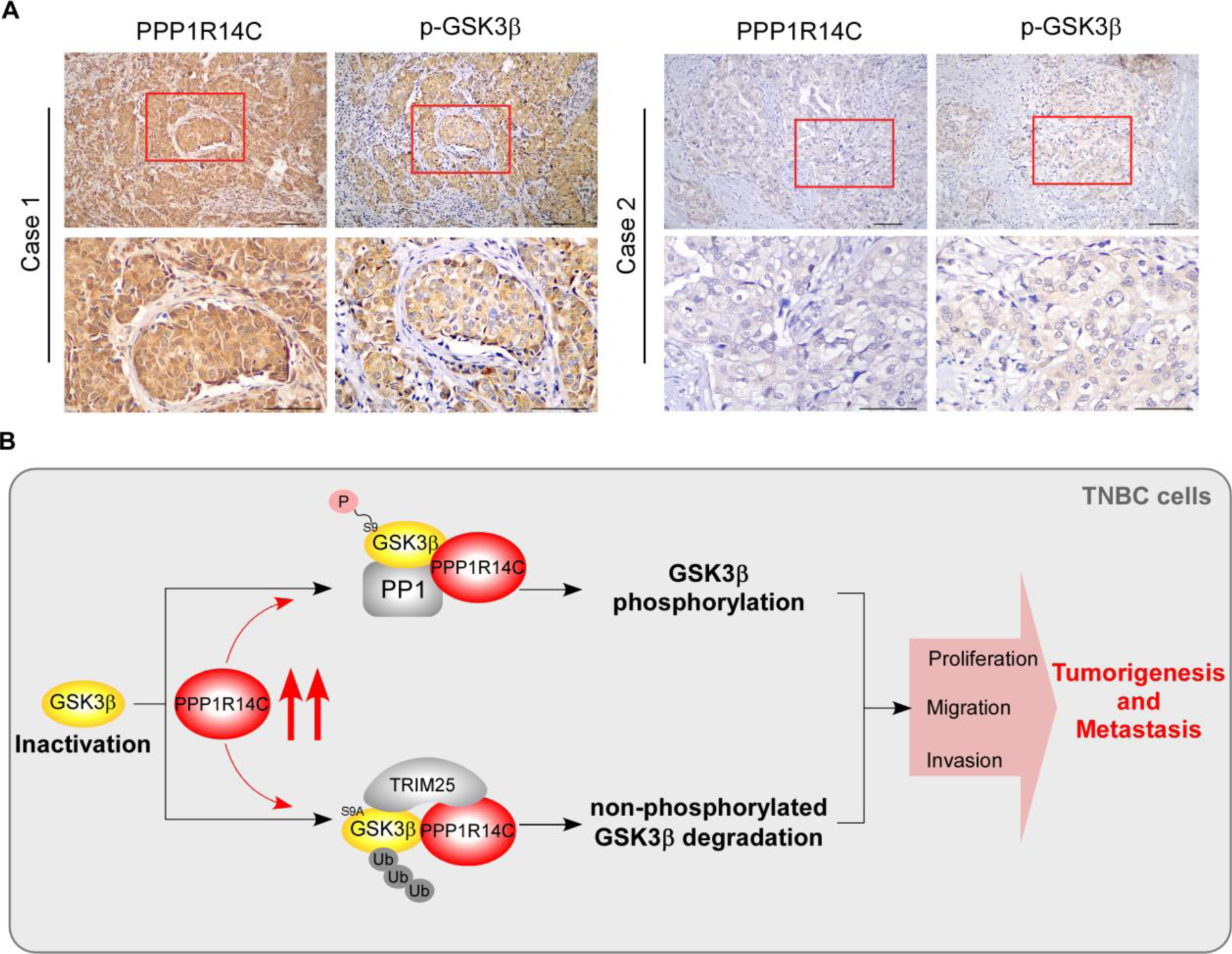
Clinical relevance of the PPP1R14C/PP1/p-GSK3β axis in TNBC. **(A)** Representative images of PPP1R14C and p-GSK3β (Ser9) IHC staining in 100 specimens from patients with breast cancer. Scale bars represent 100 μm. **(B)** Illustration of the Mechanism.

## Discussion

Patients with TNBC suffer worse prognosis than those with other subtypes because of higher rates of recurrence and limited therapeutic options. Moreover, TNBC is usually more aggressive and more likely to metastasize[35, 36]. Therefore, effective and specific regulators for TNBC are urgently needed. The present study showed that PPP1R14C was markedly overexpressed in TNBC tissues and cell lines, but not in normal tissues or non-TNBC. Furthermore, patients with TNBC and high PPP1R14C expression were more likely to have shorter 5-year OS and RFS, compared with other subtypes. PPP1R14C inactivated PP1 to sustain GSK3β phosphorylation at Ser9, which promoted the degradation of GSK3β, and contributed to the aggressiveness of TNBC. Moreover, blocking the function of PPP1R14C using C2 inhibited the malignant phenotype in TNBC cells. In conclusion, our results revealed the oncogenic role of PPP1R14C in TNBC and provided a new biomarker for effective targeted therapy.

GSK3β is implicated in many cell processes, including the regulation of transcription factors, cell-cycle progression, cell survival, apoptosis and cell migration[37]. Non-phosphorylated GSK3β is highly active in the basal-state and exerts an inhibitory effect on its downstream pathways[38]. Persistently inactive GSK3β has been found in various cancers, especially TNBC, suggesting that p-GSK3β has an oncogenic role in neoplastic disease[17, 19]. However, Cao Q et al have found that the active form of GSK3β might also function as an oncogene, because it contributed to cell proliferation by inducing S phase entry in ovarian cancer cells[18]. These apparently contradictory results suggest that the functions of GSK3β might differ according to the cell type and cellular context[37]. The present study demonstrated that PPP1R14C, together with PP1 and upstream stimulations, maintained GSK3β in a Ser9-phosphorylated and inactive state to promote tumor proliferation and metastasis in TNBC. Furthermore, PPP1R14C promoted non-phosphorylated GSK3β ubiquitin-dependent degradation, which synergistically amplified and prolonged p-GSK3β signaling in TNBC. These results represent a novel mechanism that explains how GSK3β remains inactive in TNBC and further supports the oncogenic functions of p-GSK3β.

Studies have indicated that PPP1R14C regulates protein activity dependent on its inhibitory effect on PP1, which modulates the biological activities of numerous key proteins by regulating their phosphorylation states directly[20, 39]. Dóra Dedinszki et al have reported that PPP1R14C overexpression increased the phosphorylation level of RB1 and decreased the chemosensitivity of leukemia cells to chemotherapeutic drugs[23]. However, studies on PPP1R14C in human tumors are limited. The present study revealed that PPP1R14C was specifically upregulated in TNBC and PPP1R14C overexpression correlated strongly with worse prognosis of patients with TNBC. Furthermore, PPP1R14C promoted cell proliferation, migration, invasion, tumor growth and lung metastasis of TNBC cells by sustaining the phosphorylation state of GSK3β at Ser9 and degrading non-phosphorylated GSK3β, demonstrating the novel oncogenic functions of PPP1R14C, which might serve as a targeted therapeutic strategy in TNBC. Although PPP1R14C was reported as down-regulated in breast cancer cells, we detected PPP1R14C in all the breast cancer subtypes and discovered that PPP1R14C was specifically overexpressed in TNBC, which was supported by public cancer datasets from the GEO and TCGA[40]. The discovery suggested that PPP1R14C plays a specific role in TNBC progression.

As a PP1 inhibitory protein, PPP1R14C shares an N-terminal binding domain (residues 20-24; RVFFQ) with most PP1 regulatory proteins[41]. PP1 regulatory proteins target PP1 to distinct subcellular locations or specific substrates[27, 42]. The interaction of myosin-targeting subunit M110 with PP1 enhanced its activity towards myosin P-light chains[43]. Interestingly, PPP1R14C is overexpressed in TNBC, which suggested that excessive PPP1R14C could efficiently compete for PP1 binding and disrupt PP1’s interaction with its other regulatory partners. In this study, we found that interaction of PPP1R14C with PP1 increased its specificity for p-GSK3β (Ser9), compared with other tested substrates. suggesting the PPP1R14C-PP1 interaction enhanced their affinity for p-GSK3β (Ser9) to strongly inhibit its dephosphorylation. Thus, PPP1R14C-modulated persistence of p-GSK3β might occur specifically in TNBC.

In summary, we identified an oncogenic role of PPP1R14C in TNBC in which it interacts with PP1 to maintain the phosphorylation of GSK3β at Ser9, and induces non-phosphorylated GSK3β ubiquitylation and degradation, contributing to aggressive phenotype of TNBC (Figure 8B). These findings increase our awareness of the regulatory mechanisms of GSK3β activity in TNBC, and provide new directions for therapeutic strategies to treat TNBC. Further investigation is warranted to determine whether a PPP1R14C inhibitor may be an effective approach to cure TNBC.

## Materials and Methods

### Cell lines and cell culture

Breast cancer cell lines (MCF-7, ZR75-30, SK-BR-3, HCC1937, MDA-MB-231 and BT-549) and Hela were provided by the American Type Culture Collection (Manassas, VA, USA) and the SUM159PT breast cancer cell line was from Asterand Bioscience (Detroit, MI, USA). BT-549 cells were cultured in RPMI 1640 with 10% FBS, SUM159PT was cultured in Ham’s F-12 with 5% FBS, 1 μg/ml hydrocortisone, 5 μg/ml insulin and 10 mM HEPES, and the other cells were cultured in Dulbecco’s modified Eagle’s medium (DMEM) with 10% fetal bovine serum, 100 U/ml penicillin, and 100 μg/ml streptomycin. Cells were maintained at 37°C in a 5% CO2 incubator. Cells were routinely tested for mycoplasma contamination using the Lookout Mycoplasma PCR Detection Kit (#MP0035; Sigma-Aldrich). Cells were treated with the proteasome inhibitor MG132 (ApexBio, Hsinchu, Taiwan, China) at 10 µM to inhibit proteasome-mediated degradation and with cycloheximide (ApexBio) at 50 µg/mL to inhibit translation. All cell lines were authenticated using short tandem repeat (STR) profiling before experiments.

### Patient information and tissue specimens

We collected 10 adjacent normal and 150 cancer specimens that were histopathologically diagnosed at the Sun Yat-sen University Cancer Center from 2004 to 2012. All patients eligible for this study accepted surgery and were followed-up regularly. Detailed clinicopathological data and survival data were collected (Supplementary Table S1). The study was approved by the ethics committee of Sun Yat-sen University Cancer Center and was performed according to Declaration of Helsinki.

### Quantitative real-time PCR (qPCR)

Total RNA was isolated from cells or human tissue using TRIzol (Invitrogen, Carlsbad, CA, USA) according to manufacturer’s instructions. cDNA was synthesized from total RNA (2 µg) after adding RNase-free DNase. qPCR was performed in triplicate using 1 µL of cDNA in a standard SYBR premix Ex Taq (Takara, Shiga, Japan) on the CFX96 Real-Time PCR Detection System (Bio-Rad, Hercules, CA, USA). Glyceraldehyde-3-phosphate dehydrogenase (GAPDH) served as an internal control.

### Western blotting

Total cellular proteins were extracted using Radioimmunoprecipitation assay lysis buffer (Thermo Scientific, Waltham, MA, USA) quantified using a Bio-Rad DC protein assay kit II, separated by electrophoresis on 8–15% SDS-PAGE gels and electro transferred onto a Hybond ECL transfer membrane (Amersham Pharmacia, Piscataway, NJ, USA). The membranes were blocked with 5% skimmed milk and incubated with the appropriate primary antibodies. The antigen-antibody complexes on the membrane were detected using labeled secondary antibodies and enhanced chemiluminescence reagents (Thermo Scientific). The antibodies and their dilutions were as follows: anti-PPP1R14C (#PA5-50996, dilution 1:1000, Invitrogen), anti-PP1 (#sc-7482, dilution 1:500, Santa Cruz Biotechnology, Sant Cruz, CA, USA), anti-GSK3β (#22104-1-AP, dilution 1:2000, Proteintech, Rosemont, IL, USA), anti-p-GSK3β (Ser9) (#9323, dilution 1:1000, CST, Danvers, MA, USA), anti-HA (H6908, dilution 1:1000, Sigma-Aldrich, St. Louis, MO, USA), anti-Flag (F7425, dilution 1:1000, Sigma-Aldrich), anti-MYC (05-724, dilution 1:1000, Sigma-Aldrich), anti-GAPDH (#5174, dilution 1:1000, CST), anti-p-p53 (Ser15) (MA5-15229, dilution 1:1000, Invitrogen), anti-p-CREB (Ser133/Ser63) (MA1-114, dilution 1:1000, Invitrogen), anti- p-eIF2A (Ser51) (#3398, dilution 1:1000, CST), anti-p-Aurora A (Thr288) (MA5-14904, dilution 1:1000, Invitrogen), anti-p-Slug (S246) (ab63568, dilution 1: 500, Abcam), p-c-myc (T58) (ab185655, dilution 1:1000, Abcam), anti-c-Jun (Thr239) (PA5-104748, dilution 1:1000, Invitrogen) and anti-p-SMAD3 (S423/425) (#9520, dilution 1:1000, CST) and then exposed to horseradish peroxidase (HRP)-conjugated secondary anti-mouse or rabbit antibodies. Immunoreactive proteins were detected using enhanced chemiluminescence (ECL) (Amersham Pharmacia, Little Chalfont, UK).

### Immunohistochemistry

IHC was performed to detect PPP1R14C in 10 adjacent normal and 150 breast cancer specimens using anti- PPP1R14C (#PA5-50996, Invitrogen) antibody. The staining index (SI) of tissues was evaluated using the intensity and proportion of positively stained tumor cells. Staining intensity was classified as follows: 0, no staining; 1, weak staining (light yellow); 2, moderate staining (yellow brown); 3, strong staining (brown). Scores of positively stained cell proportions were: 0, no positive; 1, < 10%; 2, 10–35%; 3, 35–75%; 4, > 75%. The levels of the indicated proteins were determined using the SI. The SI consisted of possible scores of 0, 1, 2, 3, 4, 6, 8, 9 and 12. High and low expression of PPP1R14C were defined as an SI ≥ 6 and SI < 6, respectively, according to the heterogeneity with the log-rank test statistics with respect to 5-year OS and RFS.

### Plasmids and generation of stably transfected cell lines

Transient plasmid transfection was performed using Lipofectamine 3000 (Invitrogen) according to the manufacturer’s instructions. To construct stable PPP1R14C-knockdown cells, MDA-MB-231 and SUM159PT cell lines were infected with lentiviruses containing two different short hairpin RNAs targeting PPP1R14C (GeneChem, Shanghai, China). For overexpression, the full-length PPP1R14C cDNA was identified and cloned to generate a Flag- PPP1R14C construct. To construct stable PPP1R14C-overexpressing cells, lentiviruses were constructed, infected into MDA-MB-231 and SUM159PT cells, and selected using puromycin.

### Colony formation assay

Cells (400 per plate) were cultured in 6-well plates for 14 days. The formed colonies were fixed with ethanol, stained with 1% crystal violet, and counted.

### Anchorage-independent growth ability assay

MDA-MB-231 and SUM159PT cells were trypsinized and suspended in 0.66% agar (Sigma) plus 2 ml of DMEM (with 10% fetal bovine serum) in a 6-well plate (5000 cells per well). Cells mixed with culture medium were plated above a layer containing a mixture of 1.32% agar plus medium. Colonies were counted and photographed after 10 days.

### Flow cytometry analysis

Cell cycle was analysed on a flow cytometer FACScan instrument (Beckman Coulter, Brea, CA, USA). 20000 cells were washed, fixed, pelleted, and incubated in bovine pancreatic RNAse (Sigma). The cells were then stained with propidium iodide (Sigma-Aldrich). The specific procedure was reported previously (Lin et al., 2010).

### Immunoprecipitation assay

Lysates were prepared from the indicated cancer cells using lysis buffer (150 mM NaCl, 10 mM HEPES, [pH 7.4], 1% NP-40). The lysates were incubated with protein G agarose (IP04, Millipore), Flag affinity agaros (A2220, Millipore) or HA affinity agarose (A2095, Millipore) overnight at 4° C. Beads containing affinity- bound proteins were washed six times with immunoprecipitation wash buffer (150 mM NaCl, 10 mM HEPES, [pH 7.4], and 0.1% NP-40), followed by elution with 1 M glycine [pH 3.0]. The eluates were then mixed with sample buffer, denatured, and electrophoresed for western blotting analysis.

### Xenograft tumor models

Female BALB/c-nu mice (5- 6 weeks old, 18- 20 g) were purchased and housed in barrier facilities on a 12-h light/dark cycle. The Institutional Animal Care and Use Committee of Sun Yat-sen University approved all the experimental procedures. To establish the subcutaneous xenograft model, 1 x 10^6^ SUM159PT-vector/scramble, -PPP1R14C, and -shPPP1R14C cells were injected subcutaneously into the fat pads of mice. The tumor volumes were determined every week. The tumor volume was calculated using the following equation: (L*W^2^) / 2 (L = length, W = width). The mice were sacrificed after 7 weeks, and the tumors were isolated and weighed. Serial 6.0-μm sections were cut and stained with anti- PPP1R14C and anti- Ki-67 (cat. no. ZM-0166; Zhongshanjinqiao Bio-Reagent Company) antibodies to determine the level of proliferation.

For lung colonization and spontaneous metastasis models, mice were randomly divided into groups (n = 6 per group), and intravenously or subcutaneously injected with 2 × 10^5^ MDA-MB-231 and 4T1 cells. Bioluminescence imaging of tumor colonization and growth in lung tissues were evaluated using the Xenogen IVIS spectrum imagining system (Caliper Life Sciences, Hopkinton, MA, USA) as described previously[44]. To block the PPP1R14C/PP1/p-GSK3β axis, mice were injected intraperitoneally with C2 (100 mg/kg) once per day[45]. 12 weeks later, the mice were killed, and lungs were removed and fixed in formalin and paraffin-embedded for hematoxylin and eosin staining. The number of lung metastases in each group was counted under five random low power fields and presented as the mean ± S.D..

### Statistical analysis

Statistical analyses were performed using the SPSS version 19.0 statistical software package (IBM Corp., Armonk, NY, USA). The values presented are expressed as the mean with S.D.. All experiments were performed at least in triplicate. Statistical tests for data analysis included two-tailed Student’s t test, Mann-Whitney U test and χ2 test. Kaplan-Meier methodology was used to evaluate survival probabilities and log-rank test was used to compare survival difference on univariate analysis. Multivariate statistical analysis was performed using a Cox regression model. *P* < 0.05 was considered statistically significant. No statistical method was used to predetermine sample sizes; however, our sample sizes were similar to those reported in previous studies [44].

## Acknowledgements

Not applicable.

## Author contributions

Y.J., L.K. and H.X. carried out most of the experimental work. S.L. and D.S. conducted the immunofluorescence and IHC staining analysis. M.W. and Y.L. performed the IP assays. H.X., Y.X. and M.Y. performed the western blotting and public datasets anlysis. Y.J., L.K. and Y.O. conducted the cell culture, stable cell line establishment and *in vitro* functional assays. X.H. and X.C. conducted the animal model experiments. Y.H. and X.C. collected the clinical samples and patient information. P.L. and W.W. conceived of the project, designed most of the experiments, wrote the manuscript, and supervised project.

## Supplementary Figure Legends

**Supplementary Figure 1.**
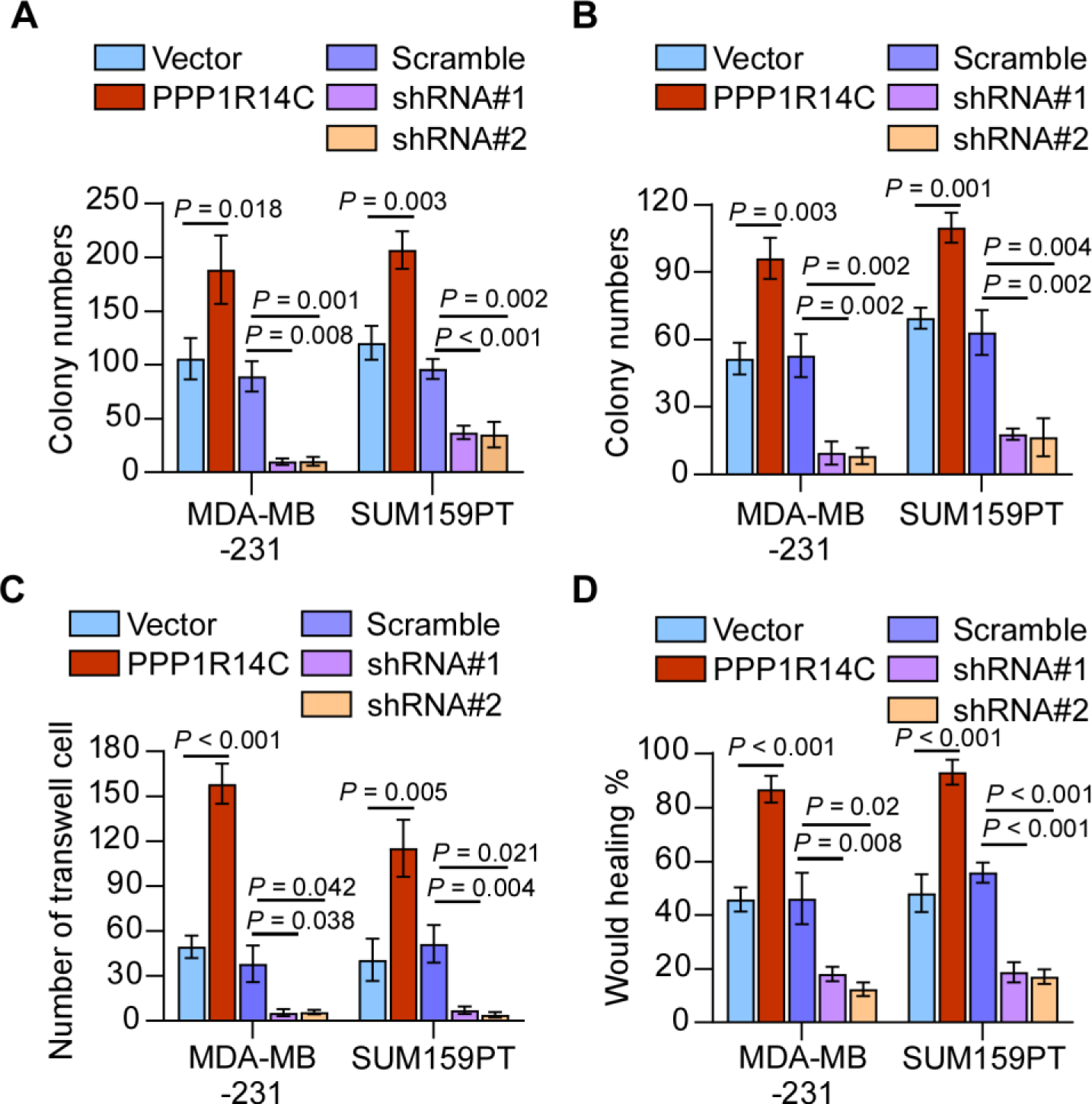
**(A-D)** The quantifications of colony formation (A), soft agar (B), transwell (C), wound healing (D) assays for the indicated cell lines. Two-tailed Student’s t test was used.

**Supplementary Figure 2.**
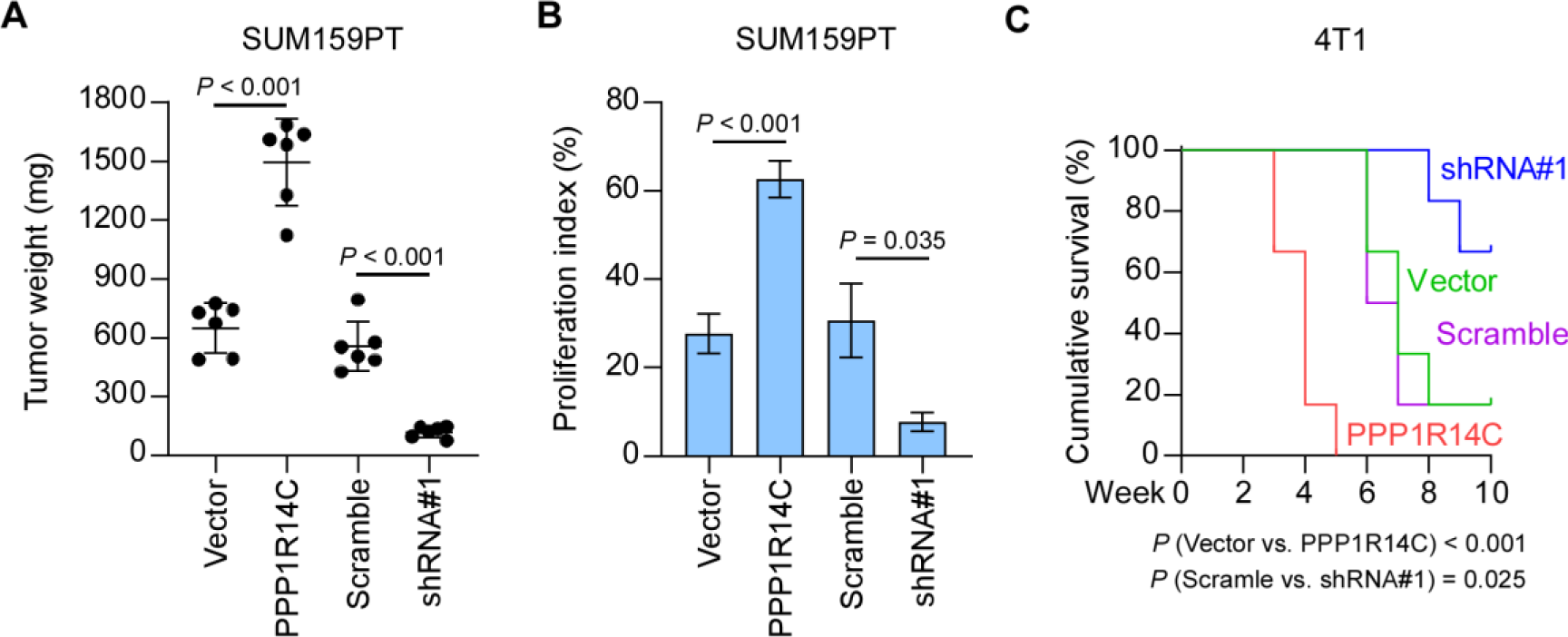
**(A)** The tumor weights in the indicated groups are measured. **(B)** Percentage of Ki-67 were shown in the indicated cells. **(C)** Kaplan–Meier survival curves of mice injected with the indicated cells.

**Supplementary Figure 3.**
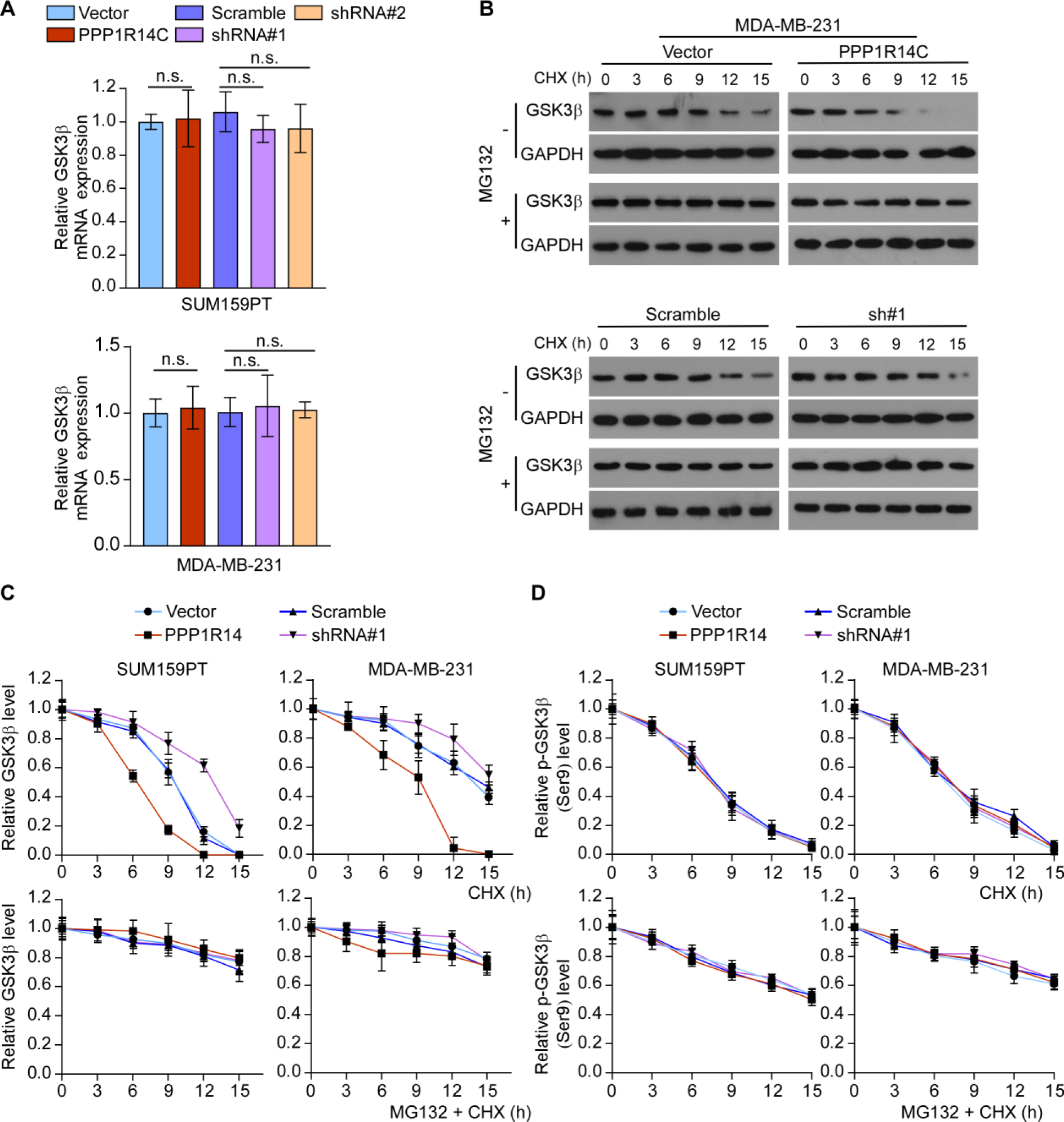
**(A)** Real-time PCR analysis of GSK3β in control, PPP1R14C-overexpressed and -knockdown TNBC cells. **(B)** Western blot analysis of GSK3β protein in MDA-MB-231 cells treated with CHX (50 µg/mL) for 0, 30, 60, or 120 min plus with or without MG132 (10 µM) treatment. GAPDH was used as loading control. **(C)** The statistical graph of Figure 6B and Supplementary Figure 4B. **(D)** The statistical graph of p-GSK3β protein in TNBC cells treated with CHX (50 µg/mL) for 0, 30, 60, or 120 min plus with or without MG132 (10 µM) treatment.

**Supplementary Figure 4.**
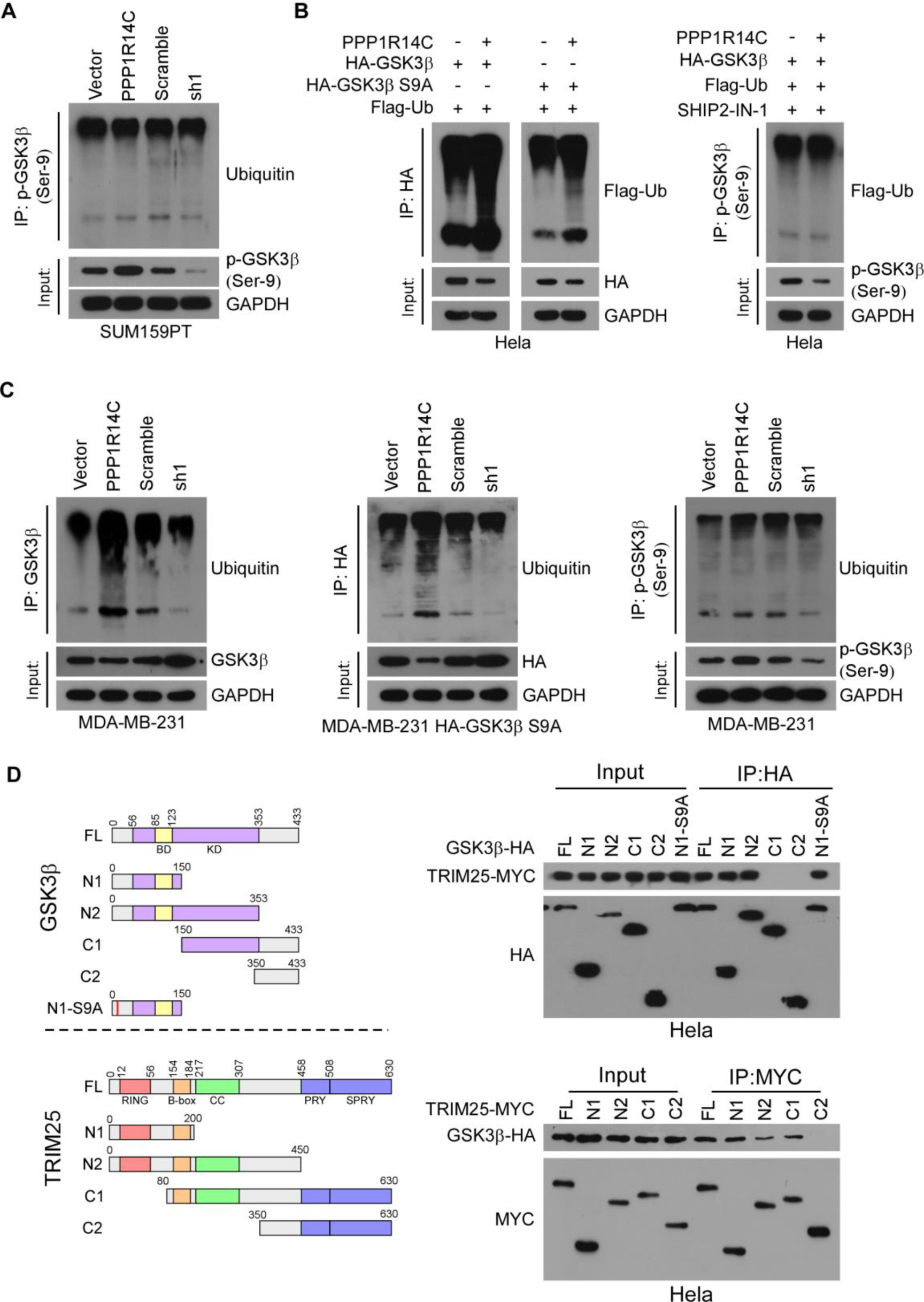
**(A)** Effect of ubiquitination on p-GSK3β by immunoprecipitation assay in SUM159PT cells using anti-p-GSK3β antibody to pull down. **(B)** Effect of ubiquitination on GSK3β, non-phosphorylated GSK3β (S9A) and p-GSK3β by immunoprecipitation assay in Hela cells using anti-HA or anti-p-GSK3β antibody to pull down, respectively. **(C)** Effect of ubiquitination on different status of GSK3β by immunoprecipitation assay in MDA-MB-231 cells. **(D)** Left panel: the truncated mutants of GSK3β and TRIM25 were based on known functional domain. BD: binding domain; KD: kinase domain; RING: RING finger domain; B-box: B-box domain; CC: coiled-coil domain; PRY/SPRY: PRY/SPRY domain. Right panel: detailed interactions between GSK3β and TRIM25 were analyzed by IP assays.

**Supplementary Figure 5.**
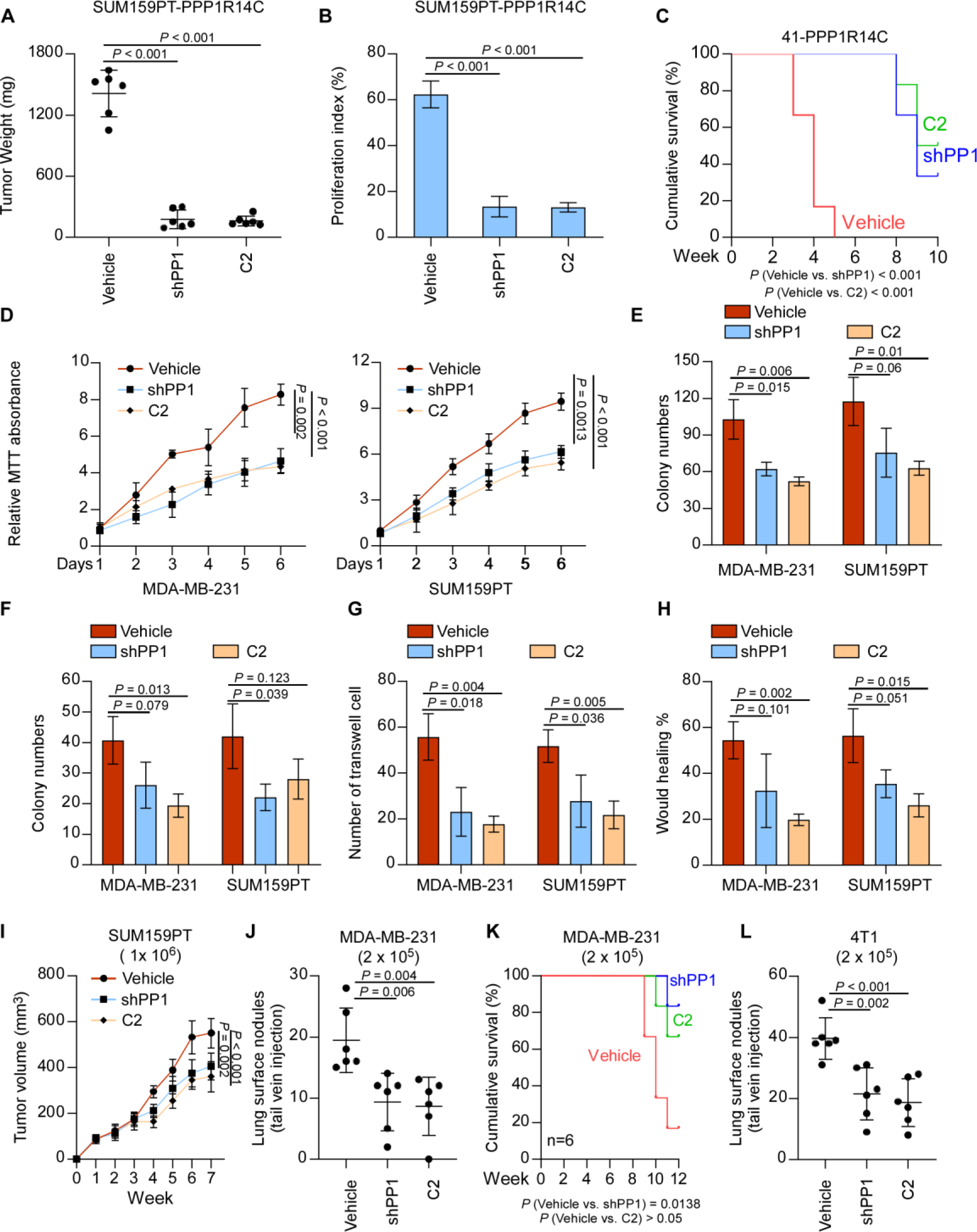
**(A-B)** The tumor weights and proliferation index in each group are shown. **(C)** Kaplan–Meier survival curves of mice. **(D-H)** MTT (D), colony formation (E), soft agar (F), transwell (G), and wound healing (H) assays were performed in vehicle, shPP1, or C2 (10 mM) in MDA-MB-231 and SUM159PT cells. Data represent the means ± S.D. of three independent experiments. Two-tailed Student’s t test was used. **(I)** The tumor volumes in MDA-MB-231-vehicle, -shPP1 and C2 treatment groups are shown. **(J)** *In vivo* metastasis assays of MDA-MB-231-vehicle, -shPP1 and C2 treatment groups. The visible surface metastatic lesions were counted. **(K)** Kaplan–Meier survival curves of mice. **(L)** The visible surface metastatic lesions of lungs were counted. Two-tailed Student’s t test and log-rank test were used.

**Supplementary Figure 6.**
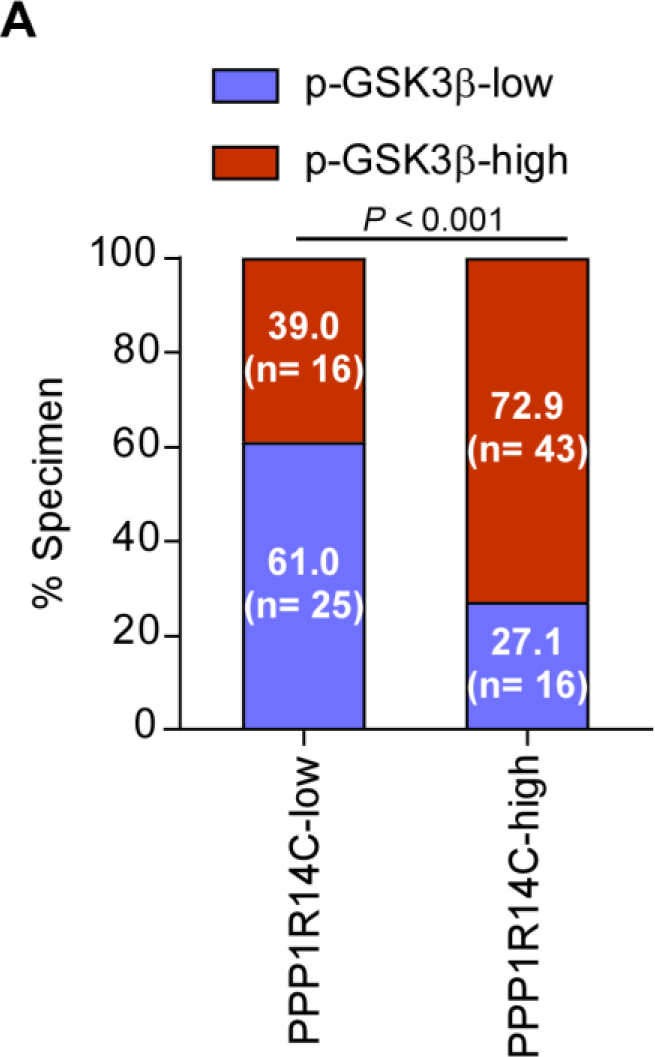
**(A)** Correlation analysis revealed that PPP1R14C expression was significantly associated with p-GSK3β (Ser9) expression in patient specimens (χ^2^ test).

